# Genomic Surveillance of *Salmonella* spp. in the Philippines, 2013-2014

**DOI:** 10.1101/2022.02.14.480363

**Authors:** Marietta L. Lagrada, Silvia Argimón, Janice B. Borlasa, Jaywardeen P. Abad, June M. Gayeta, Melissa L. Masim, Agnettah M. Olorosa, Victoria Cohen, Benjamin Jeffrey, Khalil Abudahab, Sonia B. Sia, Charmian M. Hufano, John Stelling, Matthew T.G. Holden, David M. Aanensen, Celia C. Carlos, the Philippines Antimicrobial Resistance Surveillance Program

## Abstract

Increasing antimicrobial resistance (AMR) in *Salmonella* has been observed in the Philippines. This study aims to utilize whole genome sequencing (WGS) to characterize the population and AMR mechanisms of *Salmonella* captured by the Philippine Antimicrobial Resistance Surveillance Program (ARSP) and contrast to traditional laboratory methods.

We sequenced the whole genomes of 148 *Salmonella* Typhi (*S*. Typhi) and 65 non-typhoidal *Salmonella* (NTS) collected in the Philippines in 2013–2014. From the genome sequences, we determined the serotype, multilocus sequence type, presence of determinants of antimicrobial resistance and relatedness between isolates. We also compared the genotypic predictions of serotype and AMR to the phenotypic data.

AMR rates in S. Typhi were low, with sparse acquisition of mutations associated with reduced susceptibility to fluoroquinolones or extended-spectrum beta-lactamases (ESBL) genes. In contrast, three quarters of NTS isolates were insusceptible to at least one antimicrobial, with more than half carrying mutations and/or genes linked to resistance to fluoroquinolones. ESBL genes were detected in five genomes that also carried other AMR determinants. The population of *S*. Typhi was dominated by the likely endemic genotype 3.0, which also caused of a putative local outbreak susceptible to antibiotics. The main NTS clades were global epidemic *S*. Enteritidis ST11 and the monophasic variant of *S*. Typhimurium (I 4,[5],12:i:-) ST34, which had frequently been serotyped as *S*. Typhimurium in the laboratory.

This was the first time that *Salmonella* isolated from the Philippines were characterized by WGS and we provide evidence of its utility for ongoing surveillance at the ARSP.

## INTRODUCTION

*Salmonella enterica* is a common cause of gastroenteritis and bacteraemia worldwide. ^1^ Although *S. enterica* comprises >2600 serovars, most human infections are caused by a limited number of serovars with different clinical presentations. The typhoidal *Salmonella* include *S*. Typhi and *S*. Paratyphi A, B and C, and are human-host restricted organisms that cause enteric fever, a systemic disease that disproportionally affects children in south-central and southeast Asia and sub-Saharan Africa and is treated with antibiotics. Other serovars are grouped as non-typhoidal *Salmonella* (NTS) and usually cause self-limiting gastroenteritis not requiring antimicrobial treatment. Less commonly, complicated invasive NTS infections that require antibiotic treatment are seen in specific populations, like the immunocompromised. ^2^

In the Western Pacific Region, invasive infectious disease agents account for 22% of the foodborne disease burden, with *S*. Typhi and *S*. Paratyphi A as the leading causes. Diarrhoeal disease agents account for 14% of the foodborne disease burden, with NTS the second leading cause after *Campylobacter* spp. ^3^ In the Philippines, the most common NTS serovars are *S*. Enteritidis and *S*. Typhimurium, ^4^ which parallels trends in the Western Pacific Region and worldwide. ^5, 6^

Antimicrobial resistance (AMR) in foodborne pathogens, including *S. enterica*, is a major concern for public health globally. In recent years, rising rates of fluoroquinolone and third-generation cephalosporin resistant *S. enterica* in humans have been reported. ^1^ In the Philippines, resistance rates of *S*. Typhi against first and second-line antibiotics remained below 10% since 2004, although resistance rates for nalidixic acid increased slightly starting around 2012. ^4^ In contrast, resistance rates of NTS against first and second-line antibiotics above 10% were recorded since the year 2004, with resistance to ceftriaxone (third-generation cephalosporin) ranging between 10 and 20%. Resistance to third-generation cephalosporin generally arises via the acquisition of extended-spectrum beta-lactamases (ESBL) or AmpC hydrolytic enzymes. ^1^ Resistance to fluoroquinolones such as ciprofloxacin may be due to mutations in the quinolone-resistance determining region (QRDR) of the *gyr* and *par* genes or the acquisition of plasmid-mediated quinolone resistance (PMQR) genes. ^7^

Until recently, AMR surveillance by the Philippine Department of Health Antimicrobial Resistance Surveillance Program (DOH-ARSP) had involved exclusively phenotypic methods. In this study we sequenced the whole genomes of *Salmonella* isolates collected by the ARSP between 2013 and 2014 using WGS to describe their population, identify AMR determinants, and determine the concordance between laboratory tests and genotypic predictions of serotype and resistance.

## METHODS

### Bacterial Isolates

A total of 258 *S*. Typhi and 326 NTS isolates were collected by the Philippine DOH-ARSP in 2013 and 2014 (**Table 1**), and 171 *S*. Typhi and 68 NTS isolates were referred to the ARSRL for confirmation of bacterial identification and resistance profile. Out of these, 153 *S*. Typhi and 65 NTS isolates successfully resuscitated from the biobank were submitted for whole-genome sequencing (WGS).

**Table 1.** Number of *Salmonella* isolates referred to the analysed by the Antimicrobial Resistance Surveillance Program (ARSP) and referred to the Antimicrobial Resistance Surveillance Reference Laboratory (ARSRL) during 2013 and 2014, isolates submitted for whole-genome sequencing, and high-quality genomes obtained, discriminated by sentinel site and AMR profile.

|  | S. Typhi |  |  | NTS |  |  | Grand Total |
| --- | --- | --- | --- | --- | --- | --- | --- |
|  | 2013 | 2014 | Total | 2013 | 2014 | Total |  |
| Total ARSP | 119 | 139 | 258 | 168 | 158 | 326 | 584 |
| Referred to ARSL | 84 | 87 | 171 | 31 | 37 | 68 | 239 |
| Submitted for WGS | 78 | 75 | 153 | 31 | 34 | 65 | 218 |
| High-quality genomes | 76 | 72 | 148 | 31 | 34 | 65 | 213 |
| *By sentinel site* |  |  |  |  |  |  |  |
| BGH | 2 | 3 | 5 | 2 | 2 | 4 | 9 |
| BRT | 4 | 3 | 7 | 0 | 0 | 0 | 7 |
| CMC | 15 | 13 | 28 | 0 | 1 | 1 | 29 |
| CVM | 5 | 2 | 7 | 1 | 1 | 2 | 9 |
| DMC | 3 | 3 | 6 | 1 | 4 | 5 | 11 |
| EVR | 13 | 9 | 22 | 0 | 1 | 1 | 23 |
| FEU | 2 | 1 | 3 | 0 | 2 | 2 | 5 |
| GMH | 5 | 9 | 14 | 0 | 0 | 0 | 14 |
| JLM | 0 | 0 | 0 | 0 | 4 | 4 | 4 |
| MAR | 4 | 6 | 10 | 0 | 3 | 3 | 13 |
| MMH | 1 | 0 | 1 | 0 | 0 | 0 | 1 |
| NMC | 2 | 1 | 3 | 1 | 0 | 1 | 4 |
| RMC | 0 | 0 | 0 | 2 | 0 | 2 | 2 |
| SLH | 0 | 0 | 0 | 1 | 0 | 1 | 1 |
| STU | 1 | 1 | 2 | 12 | 10 | 22 | 24 |
| VSM | 18 | 21 | 39 | 11 | 3 | 14 | 53 |
| ZMC | 1 | 0 | 1 | 0 | 3 | 3 | 4 |
| *By AMR profile* |  |  |  |  |  |  |  |
| Susceptible | 73 | 69 | 142 | 17 | 17 | 34 | 176 |
| AMP | 0 | 0 | 0 | 4 | 7 | 11 | 11 |
| SXT | 0 | 0 | 0 | 3 | 2 | 5 | 5 |
| AMP CHL | 0 | 0 | 0 | 2 | 2 | 4 | 4 |
| AMP SXT CHL | 0 | 0 | 0 | 1 | 2 | 3 | 3 |
| AMP CIP SXT CHL | 0 | 0 | 0 | 2 | 0 | 2 | 2 |
| AMP CRO | 1 | 0 | 1 | 1 | 1 | 2 | 3 |
| AMP SXT | 0 | 0 | 0 | 0 | 1 | 1 | 1 |
| AMP CIP SXT | 0 | 0 | 0 | 1 | 0 | 1 | 1 |
| AMP CIP | 0 | 0 | 0 | 0 | 1 | 1 | 1 |
| CHL | 0 | 0 | 0 | 0 | 1 | 1 | 2 |
| CIP NAL | 2 | 3 | 5 | 0 | 0 | 0 | 5 |

### Antimicrobial Susceptibility Testing (AST)

Isolates were tested for antimicrobial susceptibility to seven antimicrobial agents, ampicillin (AMP), ceftriaxone (CRO), cefotaxime(CTX), chloramphenicol (CHL), ciprofloxacin (CIP), nalidixic acid (NAL), and trimethoprim-sulfamethoxazole (SXT) with the Vitek 2 Compact automated system (bioMérieux, Marcy-l’Étoile, France) and interpretive criteria and breakpoints from the Performance Standards for Antimicrobial Susceptibility Testing (26th edition) of the Clinical and Laboratory Standards Institute (CLSI)^13^. The ESBL phenotype and insusceptibility to quinolones were confirmed using E-test™ (bioMérieux; Marcyl’Étoile, France). Multi-drug resistant (MDR) organisms were those resistant to resistant to ampicillin, chloramphenicol and trimethoprim-sulfamethoxazole. ^8^

### Serotyping

Serological serotyping was performed using Sven-Gard method for slide agglutination with antisera from Denka Seiken (Tokyo, Japan) and S&A serotest (Thailand). *Salmonella* serotypes were determined with the White-Kauffmann classification scheme. ^9^

### DNA Extraction and Whole-Genome Sequencing

Isolates were grown on Tryptic Soy Broth overnight at 35^0^C. DNA was extracted from single colonies using Wizard^®^ Genomic DNA Purification Kit (Promega). The DNA extracts were shipped to the Wellcome Sanger Institute for sequencing on the Illumina HiSeq platform (Illumina, San Diego, CA, USA) with 100-bp paired-end reads. Raw sequence data were deposited in the European Nucleotide Archive (ENA) under the study accession PRJEB17615. Individual run and sample accessions are provided through the links to Microreact projects in the figure legends.

### Bioinformatics analysis

Genome quality was evaluated based on metrics generated from assemblies, annotation files, and the alignment of the reads to the reference genome of strains 08-00436 (accession GCF_002238275.1) or CT18 (accession GCF_000195995.1), as previously described. ^10^ Annotated assemblies were produced as described in detail previously. ^11^

Evolutionary relationships between 148 *S*. Typhi isolates were inferred from single-nucleotide polymorphisms (SNPs) by mapping the paired-end reads to the reference genome of strain CT18 (accession) as described in detail previously (Argimon et al 2020). The mobile genetic elements and repetitive sequences in the genome of CT18 previously defined ^12^, ^13^ were masked in the pseudo-genome alignment with a script available at https://github.com/sanger-pathogens/remove_blocks_from_aln. Recombination regions were removed using Gubbins v2.0.0 ^14^ and the non-recombinant SNPs were used to infer a maximum-likelihood tree with RAxML v. 8.28 ^15^ based on the generalised time reversible model with the GAMMA method of correction for among-site rate variation and 500 bootstrap replications. Pairwise SNP differences between genomes were calculated from alignments of SNP positions with a script available at https://github.com/simonrharris/pairwise_difference_count.

Evolutionary relationships between 65 NTS isolates were inferred from core genome SNPs. The core genome was determined with Roary v3.12.0, ^16^ using a blastp percentage identity of 95% and a core definition of 99%. SNPs were identified in the core genome alignment with snp-sites v2.4.0 ^17^ and a tree was obtained with RAxML as described above.

Serotype ^18^ and multi-locus sequence type (MLST, ^19^) information were derived from all *Salmonella* assembly sequences with Pathogenwatch ^20^, as well as genotype information for *S*. Typhi. ^21^

Known AMR genes and mutations were identified in the *S*. Typhi assemblies using Pathogenwatch, and in the NTS genomes from sequence reads using ARIBA ^22^ and the Resfinder ^23^ (genes) and Pointfinder ^24^ (mutations) databases. The genotypic predictions of AMR (test) were compared to the phenotypic results (reference), and the concordance between the two methods was computed for seven antimicrobials. Isolates with either a resistant or an intermediate phenotype were considered non-susceptible for comparison purposes.

To contextualize the *S*. Typhi genomes, we compared to global genomes belonging to genotypes 3.0 (n=51), 3.2.1 (n=70), and 4.1 (n=141) available on Pathogenwatch (as of May 2021), which clusters the genomes based on genetic similarity as described in detail previously. ^20^

## RESULTS

### Demographic and Clinical Characteristics of the *Salmonella* Isolates

Out of the 218 *Salmonella* isolates sequenced, 5 were excluded based on genome quality (**Table 1**). The demographic and clinical characteristics of the remaining 213 isolates (148 *S*. Typhi, 65 NTS) are summarized on **Table 2**. The majority of the patients were male overall (126/213, 59.2%), but a pronounced difference in the distribution of patient sex was observed for NTS (64.6% male, 35.4% female). The group aged 0-14 years old had the highest percentage of *S*. Typhi (60.1%, 89/148) and NTS (47.7%, 31/65) infections. NTS infections were also frequent in patients 45-80 years old (38.5%, 25/65), while *S*. Typhi infections were rare in this age group (4.1%, 6/148). The vast majority of the *S*. Typhi isolates were from blood (137/148, 92.6%), while the NTS isolates were recovered from blood and stool in similar proportions (25/65 or 38.5% and 22/65 or 33.8%, respectively).

**Table 2.** Demographic and clinical characteristics of *Salmonella* culture-positive patients with genomes included in this study (n=213).

| Characteristic |  | Number of Isolates |  |
| --- | --- | --- | --- |
|  |  | SAT | NTS |
| Sex | Male | 84 | 42 |
|  | Female | 64 | 23 |
| Age (in years) | <1 | 2 | 11 |
|  | 1-4 | 16 | 14 |
|  | 5-14 | 71 | 6 |
|  | 15-24 | 26 | 3 |
|  | 25-34 | 20 | 4 |
|  | 35-44 | 4 | 0 |
|  | 45-54 | 3 | 9 |
|  | 55-64 | 2 | 9 |
|  | 65-80 | 1 | 7 |
|  | >81 | 0 | 1 |
|  | Unknown | 3 | 1 |
| Patient type | Inpatient | 133 | 58 |
|  | Outpatient | 14 | 7 |
|  | Unknown | 1 | 0 |
| Specimen Type | Abscess | 1 | 2 |
|  | Aspirate | 0 | 2 |
|  | Blood<sup>a</sup> | 137 | 25 |
|  | Cerebrospinal fluid<sup>a</sup> | 0 | 3 |
|  | Stool | 7 | 22 |
|  | Urine | 3 | 0 |
|  | Fluid | 0 | 1 |
|  | Tracheal aspirate | 0 | 1 |
|  | Wound | 0 | 9 |

<sup>a</sup> Invasive specimen.

### Concordance Between Phenotypic and Genotypic serotyping and AMR

We determined the serotype of *Salmonella* organisms both by serological methods and genoserotyping. We predicted *S*. Typhi only among typhoidal *Salmonella* and fifteen different serotypes among the NTS, with *S*. Enteritidis (*n*=21), and monophasic variant I 4,[5],12:i:-of *S*. Typhimurium (*n*=16) being the most frequent. The concordance between genoserotyping and serological serotyping was 91.1% overall (194/213), 100% for typhoidal *Salmonella* (148/148 *S*. Typhi) and 70.8% for NTS (46/65). Genoserotyping predicted the monophasic *S*. Typhimurium serovar (I 4,[5],12:i:-) for 16 isolates serotyped in the laboratory as either *S*. Typhimurium (antigenic formula 1,4,[5],12:i:1,2, *n*=12) or Group B O:4;12;i;- (*n*=4, **Figure 1A**). In addition, genoserotyping predicted serovars *S*. Kentucky, *S*. Virchow and *S*. Enteritidis for three isolates reported as *S*. Anatum, *S*. Javiana and *S*. Heidelberg, respectively. *S*. Enteritidis was relatively more frequent than *S*. I 4,[5],12:i:-in invasive isolates (39.9% vs 17.9%, *n*=28), while their frequencies were comparable in non-invasive isolates (27.0% vs 29.7%, *n*=37).

**Figure 1.**
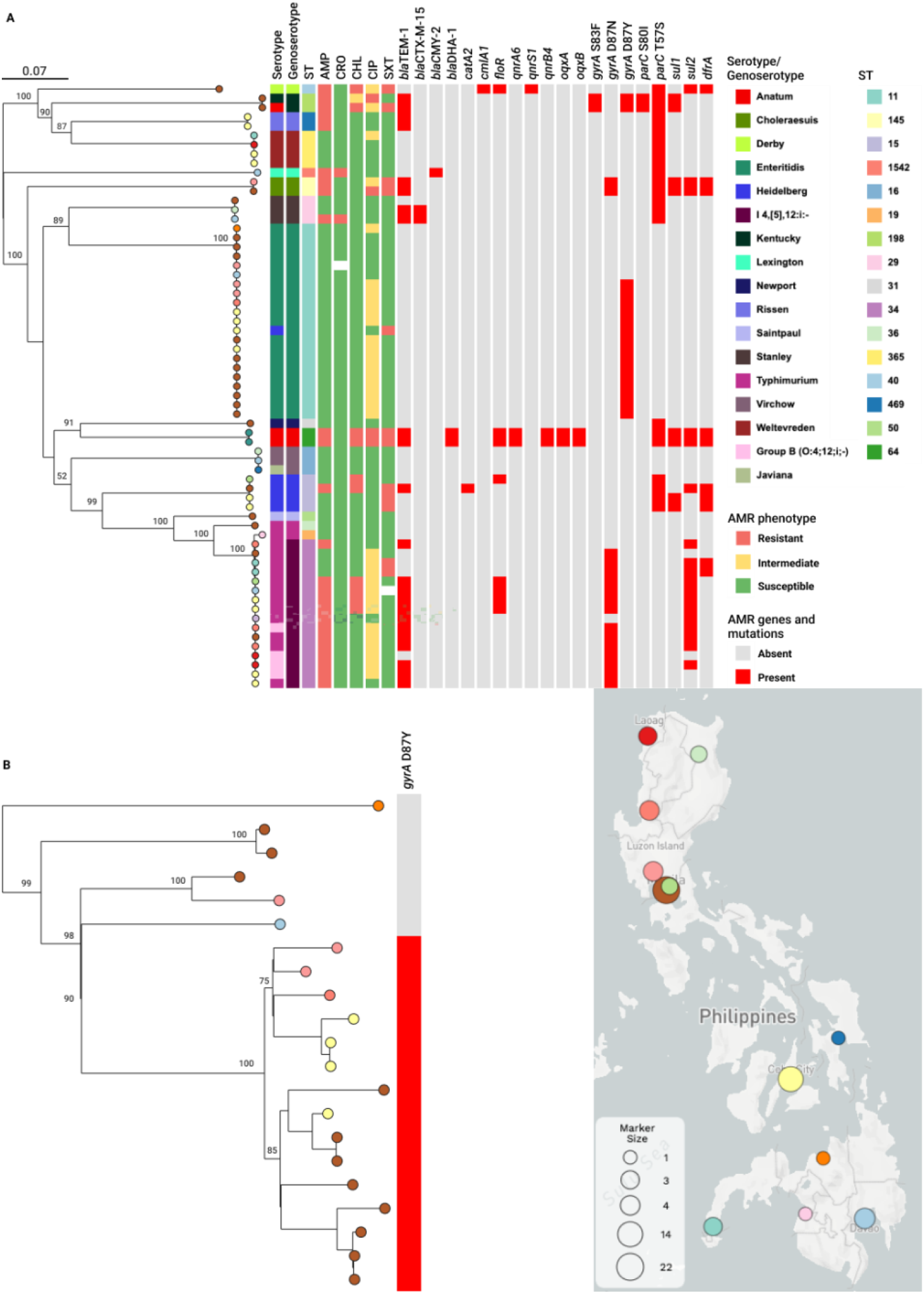
Genomic surveillance of NTS from the Philippines, 2013-2014. **A)** Phylogenetic tree of 65 isolates inferred from an alignment of 117,371 core genome SNP sites. **B)** Subtree of 21 *S*. Enteritidis isolates. The tree leaves are coloured by sentinel site as indicated on the map. The trees are annotated with bootstrap values and the tree blocks indicate the distribution of the serological serotype, genoserotype, sequence types (ST), resistance phenotype for five antibiotics and acquired resistance genes and mutations. AMP: ampicillin; CRO: ceftriaxone; CHL: chloramphenicol; CIP: ciprofloxacin; SXT: sulphamethoxazole-trimetoprim. The data are available at https://microreact.org/project/k2BC6hsaxYr1Eo5U9v71iJ-arspnts2013-2014

We also determined the susceptibilities of *Salmonella* isolates to antimicrobials (**Table 3**). *S*. Typhi isolates were largely susceptible to five antimicrobials tested. Five isolates presented both decreased susceptibility to ciprofloxacin and resistance to nalidixic acid, explained by the presence of mutations in the quinolone resistance determining region (QRDR) of the *gyrA* gene (D87N, *n*=3, and D87G, *n*=2). One isolate was resistant to ampicillin and third-generation cephalosporins (ceftriaxone and cefotaxime), mediated by the presence of the ESBL gene *bla*_CTX-M-15_ (**Table 3**). The overall concordance between phenotypic and genotypic resistance was 100% for *S*. Typhi.

**Table 3.**
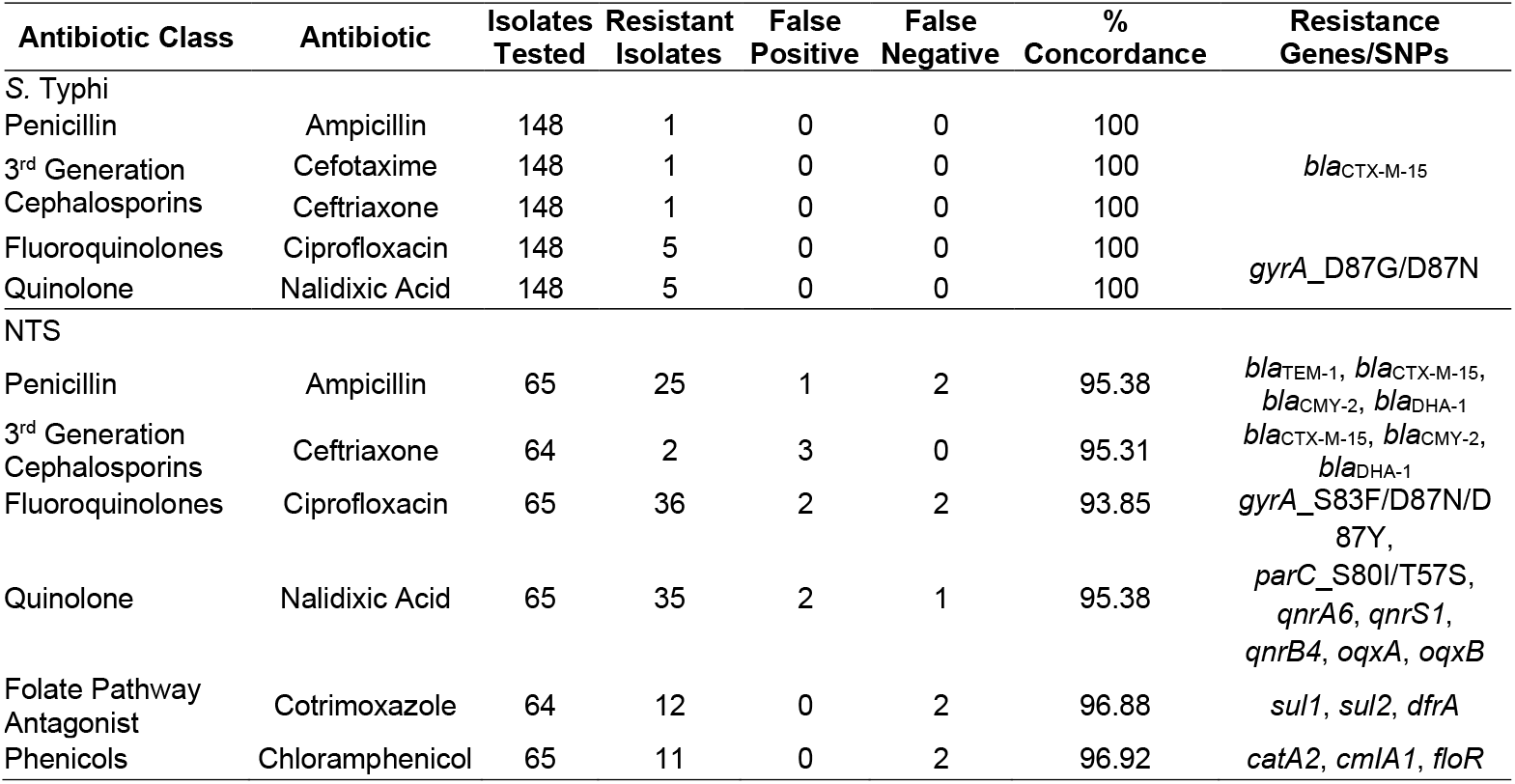
Comparison between antimicrobial susceptibility testing results and genotypic resistance for 213 *Salmonella* isolates.

The majority of NTS isolates (73.8%, 48/65) were insusceptible to at least one antimicrobial tested, most commonly to ciprofloxacin (55.4%, 36/65), and ampicillin (38.4%, 25/65). Only five isolates were MDR. Of note, the two *S*. Anatum isolates carried resistance determinants to beta-lactams (*bla*_TEM-1_, *bla*_DHA-1_), chloramphenicol (*cmlA, floR*), trimethoprim-sulfamethoxazole (*sul1, sul2, dfrA1*), ciprofloxacin (*qnrS1, qnrB4, oqxA, oqxB*, and mutation T57S in the *parC* gene), and other antibiotics not tested in the lab (*aad2, strA-strB, tet(A), mphA*, and *lnu(F)*). The overall concordance between phenotypic and genotypic resistance was 95.62% for NTS. Chloramphenicol and trimethoprim-sulfamethoxazole exhibited the highest concordances (96.92% and 96.88%, respectively). The concordance for ceftriaxone was 95.31%, and the discordance was due to three false positive results. We identified genes known to confer resistance to third-generation cephalosporins in five genomes, which also carried at least one other AMR determinant. The ESBL gene *bla*_CYM-2_ gene was found in the only *S*. Lexington isolate, the ESBL gene *bla*_CTX-M-15_ was identified in two *S*. Stanley isolates, only one of which was resistant to ceftriaxone, and the and AmpC gene *bla*_DHA-1_ was found in the two *S*. Anatum isolates, both of which were susceptible to ceftriaxone. This could be due to low expression of the inducible *bla*_DHA-1_ gene. ^25^

### In silico *genotyping*

Multi-locus sequence type and genotype were also derived from the whole-genome sequences. *S*. Typhi isolates were assigned to ST1 (132/148), ST2 (14/148) and ST5215 (2/148), and to genotypes 3.0 (121/148), 3.2.1 (14/148), 3.4 (2/148) and 4.1 (11/148). Sixteen different STs were identified among the NTS isolates and they strongly correlated to genoserotypes (**Figure 1A**), which supports *in silico* serotype assignments. Consequently, ST11 (21/65, *S*. Enteritidis) and ST34 (16/65, I 4,[5],12:i:-) were the most prevalent. *S*. Typhimurium isolates were assigned to ST19 (*n*=1) and ST36 (*n*=1).

Genotype 3.0 was found in all fourteen sentinel sites that referred *S*. Typhi isolates, while genotypes 3.4, 3.2.1 and in particular 4.1 showed more regional distributions (**Table 4** and **Figure 2A**). *S*. Enteritidis (ST11) and monophasic *S*. Typhimurium (ST34) also showed broad geographic distribution in all three island groups (Luzon in the north, Visayas in the center, and Mindanao in the south of the Philippines, **Table 4** and **Figure 1A**).

**Figure 2.**
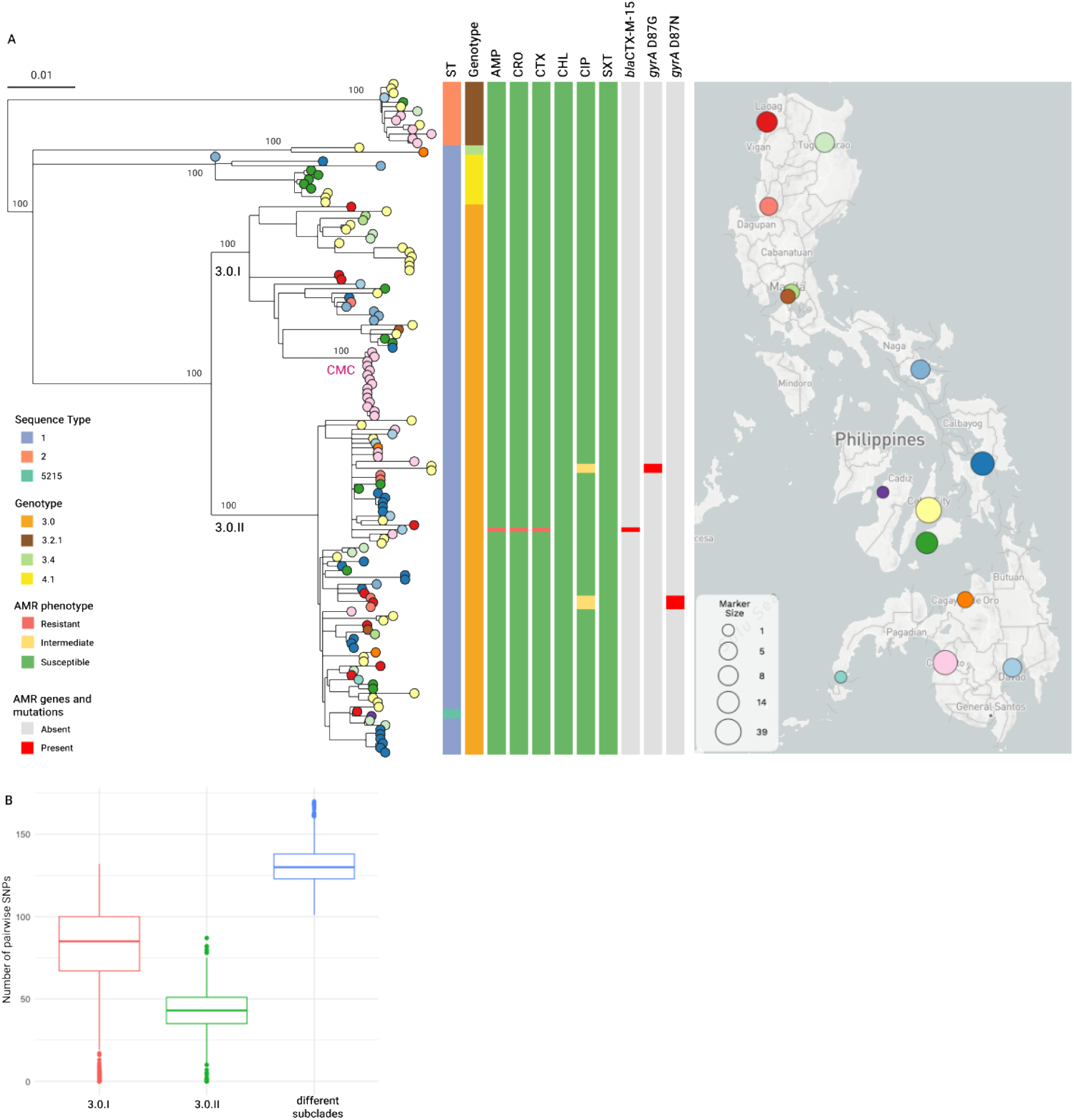
Genomic surveillance of *S*. Typhi from the Philippines, 2013-2014. **A)** Phylogenetic tree of 148 isolates inferred from an alignment of 2094 SNP sites obtained after mapping the genome sequences to the complete genome of reference strain CT18 and masking regions of mobile genetic elements and recombination. The tree leaves are coloured by sentinel site and indicated on the map. The tree is annotated with subclades within genotype 3.0 (3.0.I and 3.0.II), a putative outbreak cluster (CMC), and bootstrap values on major branches. The tree blocks indicate the distribution of the sequence types (ST), genotype, resistance phenotype for six antibiotics and acquired resistance genes and mutations. AMP: ampicillin; CRO: ceftriaxone; CTX: cefixime; CHL: chloramphenicol; CIP: ciprofloxacin; SXT: sulphamethoxazole-trimetoprim. The data are available at https://microreact.org/project/kRW7Z2TLg3FEM7rmq8sZ1e **B)** Boxplot showing the distribution of the SNP differences between pairs of genomes from genotype 3.0 belonging both to subclade 3.0.I (red), both to subclade 3.0.II (green) or one to each subclade (blue). The horizontal line indicates the median and the box indicates the interquartile range.

**Table 4.** Distribution of sequence types (ST), genoserotype and resistance profiles of *Salmonella* across the 17 sentinel sites that referred isolates. Numbers in parenthesis indicate number of isolates.

| Site<sup>a</sup> | No. of S. Typhi isolates | Prevalent ST | No. of genotypes | Prevalent genotype | AMR Resistance Profiles | No. of NTS isolates | Prevalent ST<sup>b</sup> | No. of serotypes | Prevalent serotype<sup>b</sup> | AMR Resistance Profiles<sup>c</sup> |
| --- | --- | --- | --- | --- | --- | --- | --- | --- | --- | --- |
| BGH | 5 | 1 (5) | 1 | 3.0 (5) | Susceptible (3)<br>CIP, NAL (2) | 4 | 34 (3) | 2 | I 4,[5],12:i:- (3) | AMP (3) |
| BRT | 7 | 1 (7) | 2 | 3.0 (5) | Susceptible (7) | 0 | NA | NA | NA | NA |
| CMC | 27 | 1 (21) | 2 | 3.0 (21) | Susceptible (27) | 1 | 19 (1) | 1 | SAM (1) | Susceptible (1) |
| CVM | 8 | 1 (7) | 2 | 3.0 (7) | Susceptible (8) | 2 | 16 (1) | 2 | SVR (1) | Susceptible (1) |
| DMC | 6 | 1 (5) | 2 | 3.0 (5) | Susceptible (5)<br>AMP, CRO (1) | 5 | 11 (1) | 5 | SEN (1) | Susceptible (1) |
| EVR | 22 | 1 (22) | 2 | 3.0 (21) | Susceptible (22) | 1 | 16 (1) | 1 | SVR (1) | Susceptible (1) |
| FEU | 3 | 1 (3) | 1 | 3.0 (3) | Susceptible (3) | 2 | 34 (1) | 2 | I 4,[5],12:i:- (1) | AMP, CHL (1) |
| GMH | 14 | 1 (13) | 3 | 3.0 (8) | Susceptible (14) | 0 | NA | NA | NA | NA |
| JLM | 0 | NA | NA | NA |  | 4 | 11 (3) | 2 | SEN (3) | Susceptible (3) |
| MAR | 10 | 1 (9) | 1 | 3.0 (10) | Susceptible (9)<br>CIP, NAL (1) | 3 | 34 (2) | 2 | I 4,[5],12:i:- (2) | Susceptible (1)<br>AMP (2) |
| MMH | 1 | 5215 | 1 | 3.0 (1) | Susceptible (1) | 0 | NA | NA | NA | NA |
| NMC | 3 | 1 (3) | 2 | 3.0 (2) | Susceptible (3) | 1 | 11 (1) | 1 | SEN (1) | Susceptible (1) |
| RMC | 0 | NA | NA | NA | NA | 2 | 64 (2) | 2 | SLA (2) | AMP CIP SXT CHL (2) |
| SLH | 0 | NA | NA | NA | NA | 1 | 34 (1) | 1 | I 4,[5],12:i:- (1) | AMP (1) |
| STU | 2 | 1 (2) | 1 | 3.0 (2) | Susceptible (2) | 22 | 11 (11) | 10 | SEN (11) | Susceptible (11) |
| VSM | 39 | 1 (34) | 4 | 3.0 (30) | Susceptible (37)<br>CIP, NAL (2) | 14 | 11 (4) | 5 | SEN (4) | Susceptible (3)<br>SXT (1) |
| ZMC | 1 | 1 | 1 | 3.0 (1) | Susceptible (1) | 3 | 34 (2) | 2 | I 4,[5],12:i:- (2) | SXT (2) |

<sup>a</sup> BGH: Baguio General Hospital and Medical Center; BRT: Bicol Regional Training & Teaching Hospital; CMC: Cotabato Regional Hospital and Medical Center; CVM: Cagayan Valley Medical Center; DMC: Southern Philippines Medical Center; EVR: Eastern Visayas Regional Medical Center; FEU: Far Eastern University Hospital; GMH: Governor Celestino Gallares Memorial Hospital; JLM: Jose B. Lingad Memorial Regional Hospital; MAR: Mariano Marcos Memorial Hospital and Medical Center; MMH: Corazon Locsin Montelibano Memorial Regional Hospital; NMC: Northern Mindanao Medical Center; RMC:
Rizal Medical Center; SLH: San Lazaro Hospital; STU: University of Sto. Tomas Hospital; VSM: Vicente Sotto Memorial Medical Center; ZMC: Zamboanga City Medical Center.
<sup>b</sup> For simplicity, if two or more STs/serotypes were equally prevalent at a specific site, the most prevalent of the STs/serotypes across the entire study is listed. SLA: *Salmonella* Anatum; SEN: *Salmonella* Enteritidis; SAM: *Salmonella* Typhimurium; Monophasic variant of SAM: I 4,[5],12:i:-; SVR: *Salmonella* Virchow: <sup>c</sup> The resistance profile of the prevalent ST/serotype is listed. NA: not applicable

### *Population structure of* Salmonella *in the Philippines*

The phylogenetic tree of 148 *S*. Typhi genomes was composed of four deep-branching clades that paralleled the genotype calls. However, we observed substantial diversification within the dominant genotype 3.0, which was broadly divided into two major subclades (I and II) in the tree composed by 47 and 74 genomes, respectively. The tree topology and the distribution of pairwise SNPs between genomes showed that the organisms in subclade II were genetically similar (**Figure 2A** and **B)**. Pairs of genomes belonging to subclade II were separated by median of 43 SNPs (IQR=35-51), while pairs in subclade I diverged by a median of 85 SNPs (IQR=67-100), and pairs of genomes belonging to different subclades diverged by a median of 130 SNPs (IQR=123-138). Nevertheless, both subclades were found in all three island groups. Within the more diverse subclade I, we observed a group of fifteen tightly clustered isolates from Cotabato Regional and Medical Center (CMC) recovered between May 2013 and July 2014 (**Figure 2A**). The genomes in this cluster diverged by a median of 3 pairwise SNPs (range 0-8) and carried no known resistance determinants, suggesting an outbreak of enteric fever caused by a pansusceptible strain in the population served by this hospital.

NTS isolates belonging to the same genoserotype clustered tightly together on long branches of the phylogenetic tree, thus supporting the genomic predictions. A closer inspection of the *S*. Enteritidis subtree showed that the fifteen genomes carrying mutation GyrA_D87Y associated with reduced susceptibility to ciprofloxacin formed a discreet, well-supported cluster of broader geographical distribution (**Figure 1B**). The remaining six *S*. Enteritidis genomes without any known acquired resistance determinants were found on four different branches of the subtree with narrow geographical distribution. While the distribution of invasive isolates did not significantly associate with the presence of GyrA_D87Y (*p* > 0.05), we found relatively more invasive isolates within this successful clone (9/15) than among those without the mutation (2/6).

### S. *Typhi from the Philippines in global context*

The *S*. Typhi genomes from this study were compared to global genomes from genotypes 3.0, 3.2.1, and 4.1 (**Figure 3**) available on Pathogenwatch. The Philippine genomes clustered together within each of the three genotypes and were related to genomes from countries in S or SE Asia. Surprisingly, a group of eight genomes from in Nigeria (2009-2013) also showed close relationships to the Philippine genomes within genotype 4.1, separated by between 55 and 139 SNP differences. Genotype 4.1 is widespread in both Africa and S-SE Asia, but uneven sampling of global isolates curtails our ability to establish sound transmission routes. A small number of genomes from countries in Western Europe (2007-2015) were found interspersed with Philippine genomes from genotypes 3.0 (n=5) and 3.2.1 (n=3). The epidemiological data available confirmed a travel link to the Philippines for two genomes within each genotype.

**Figure 3.**
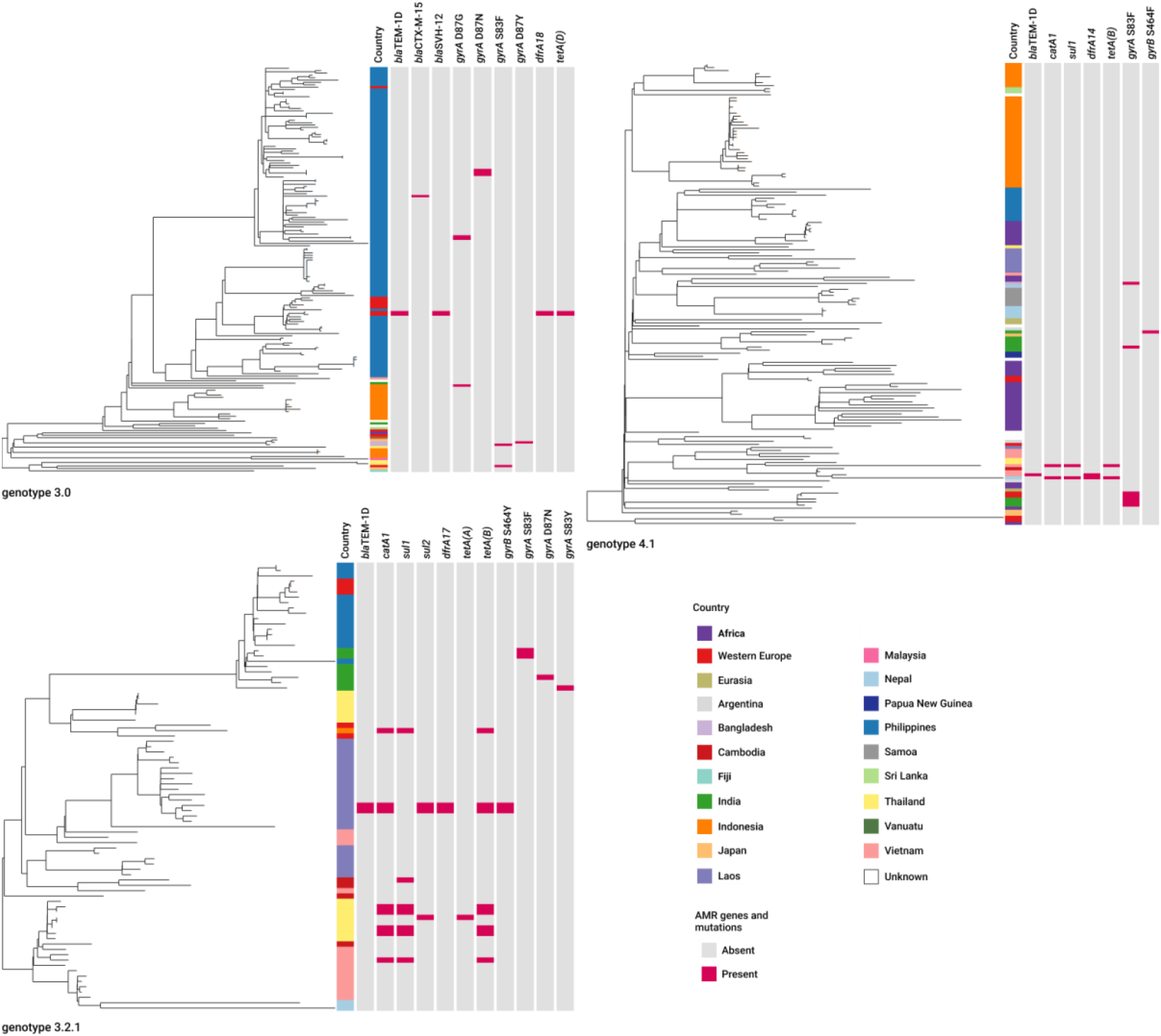
*S*. Typhi from the Philippines in global context. Phylogenetic trees of genomes belonging to genotypes 3.0, 3.2.1 and 4.1 from the Philippines and nineteen other countries or regions, generated with Pathogenwatch. Genomes from countries sparsely represented but belonging to the same continent/region were grouped to simplify the tree annotation. The trees are also annotated with the distribution of resistance determinants identified by Pathogenwatch. The data are available at https://microreact.org/project/rym1Shfy7, https://microreact.org/project/i5GByUGqNuLR9sGdDRH5hA-global-sat-321, and https://microreact.org/project/pDqxJCq7YzYy6ibxEZ2Rgk-global-sat-41

## DISCUSSION

Our study provided new insights into the *Salmonella* population from the Philippines, with important ramifications for surveillance. *Salmonella* serotyping is routinely performed at the ARSRL and it is useful for epidemiological investigations, but the serotyping scheme comprises over 2500 serovars. The genoserotyping results were largely concordant with the serological serotyping results, and confirmed that the typhoidal and non-typhoidal serovars were accurately discriminated, which is critical for patient management as typhoid fever requires antibiotic treatment. A high concordance (>94%) was reported by larger studies of *Salmonella* combining genoserotyping with MLST information. ^18, 26^ Our study also revealed inaccuracies in the serological serotyping of NTS at ARSRL, notably that most isolates typed as *S*. Typhimurium in the laboratory belonged to the monophasic variant *S*. I 4, [5],12:i:-. MLST information and phylogenetic clustering supported this assignment and highlighted the utility of the genome data.

Overall, the NTS population captured by the ARSP was diverse, with 16 clones defined by serotype and ST. A limitation of our study was that the samples available for retrospective analysis were those referred by sentinel sites to the reference lab without a consistent sampling strategy across sites. However, the serotype or ST was not contemplated for sample referral and thus our results should be representative of the population. Twelve of the NTS serotypes identified in this study, including the dominant *S*. Enteritidis and monophasic *S*. I 4, [5],12:i:-, were previously reported from retail meat in the Philippines, ^27^ suggesting a potential food-chain reservoir. The monophasic variant of *S*. Typhimurium (serovar I 4,[5],12:i:-) ST34 rose in prevalence in Europe since the early 2000s and disseminated across the world likely via the food-chain, especially pigs and pig meat, ^28^ which is the most consumed livestock meat in the Philippines. The low prevalence of MDR *Salmonella* during the survey period is in line with the absence of epidemic MDR *S*. Typhimurium clones, notably, ST313 which is dominant in sub-Saharan Africa, and the biphasic *S*. Typhimurium ST34 clone reported in Vietnam in association with HIV infection. ^29^ The combination of phylogenetic information and AMR mechanisms extracted from whole genomes led to the identification of a successful lineage of *S*. Enteritidis ST11 carrying mutation GyrA D87Y circulating across the Philippines. The relative genetic uniformity displayed *S*. Enteritidis has challenged epidemiological studies based on conventional subtyping methods ^30^ and our finding highlights the utility of the genomic data for surveillance in the Philippines beyond the resolution afforded by serotype and MLST. The *S*. Anatum organisms from this study carried the same repertoire of AMR determinants as those reported to have cause a dramatic increase of *S*. Anatum infections in Taiwan during 2016-2017. ^31^ The significance of these findings for public health grant future, more-detailed investigations into these NTS serovars and clones.

The population snapshot of *S*. Typhi showed limited diversity and predominance of genotype 3.0. The relationship between Philippine and global genomes and the diversification within this genotype suggests that this is a clone of local and persistent circulation. A limitation of our study in this respect is that the sampling encompassed only two years. We found that AMR was rare in *S*. Typhi and, in agreement with this, the genotypes found in our dataset are not known to be associated with the dissemination of single or multiple resistance, ^21^ unlike genotype 4.3.1 (haplotype H58) which was absent in our dataset. However, we observed the sporadic acquisition of resistance, notably of ESBL genes, which had only been reported before from isolates with travel history to the Philippines. ^32^ Our genomic analysis also showed evidence of a local, persistent outbreak of pansusceptible *S*. Typhi, underscoring the impact of this pathogen even in the absence of resistance.

WGS is currently being utilized for *Salmonella* surveillance in reference laboratories and international networks, and has displaced laboratory methods for both ongoing surveillance and outbreak investigations. ^33–36^ The ARSRL has implemented WGS locally but its routine use continues to be challenged in the setting of a lower middle-income economy. This first study of its utility for *Salmonella* surveillance in the Philippines supports continued application.

## ACKNOWLEDGEMENTS

This work was supported by a Newton Fund award from the Medical Research Council (UK) MR/N019296/1 and the Philippine Council for Health Research and Development. This work was also partially supported by research grant U01CA207167 from the U.S. National Institutes of Health. The contents are solely the responsibility of the authors and do not necessarily represent the official views of the funders. The funders had no role in study design, data collection and analysis, or decision to publish, or preparation of the manuscript. S.A. and D.M.A. were additionally supported by the National Institute for Health Research (UK) Global Health Research Unit on genomic Surveillance of AMR (16_136_111) and by the Centre for Genomic Pathogen Surveillance.

## REFERENCES

1. Crump JA, Sjolund-Karlsson M, Gordon MA, Parry CM. Epidemiology, Clinical Presentation, Laboratory Diagnosis, Antimicrobial Resistance, and Antimicrobial Management of Invasive Salmonella Infections. Clin Microbiol Rev. 2015;28(4):901–37.

2. Phu Huong Lan N, Le Thi Phuong T, Nguyen Huu H, Thuy L, Mather AE, Park SE, et al. Invasive Non-typhoidal Salmonella Infections in Asia: Clinical Observations, Disease Outcome and Dominant Serovars from an Infectious Disease Hospital in Vietnam. PLoS Negl Trop Dis. 2016;10(8):e0004857.

3. World Health Organization. WHO Estimates of the global burden of foodborne diseases: foodborne disease burden epidemiology reference group 2007-2015. Geneva, Switzerland; 2015.

4. Sia SB, Lagrada ML, Olorosa AM, Limas MT, Jamoralín Jr MC, Macaranas PK, et al. A Fifteen-year Report of Serotype Distribution and Antimicrobial Resistance of Salmonella in the Philippines. Philippine Journal of Pathology. 2020;5(1):19–29.

5. Galanis E, Lo Fo Wong DM, Patrick ME, Binsztein N, Cieslik A, Chalermchikit T, et al. Web-based surveillance and global Salmonella distribution, 2000-2002. Emerg Infect Dis. 2006;12(3):381–8.

6. Herikstad H, Motarjemi Y, Tauxe RV. Salmonella surveillance: a global survey of public health serotyping. Epidemiol Infect. 2002;129(1):1–8.

7. Cuypers WL, Jacobs J, Wong V, Klemm EJ, Deborggraeve S, Van Puyvelde S. Fluoroquinolone resistance in Salmonella: insights by whole-genome sequencing. Microb Genom. 2018;4(7).

8. Parry CM, Threlfall EJ. Antimicrobial resistance in typhoidal and nontyphoidal salmonellae. Curr Opin Infect Dis. 2008;21(5):531–8.

9. Grimont P, Weill F-X. Antigenic Formulae of the Salmonella serovars, (9th ed.) Paris: WHO Collaborating Centre for Reference and Research on Salmonella. Institute Pasteur. 2007:1–166.

10. Argimon S, Masim MAL, Gayeta JM, Lagrada ML, Macaranas PKV, Cohen V, et al. Integrating whole-genome sequencing within the National Antimicrobial Resistance Surveillance Program in the Philippines. Nat Commun. 2020;11(1):2719.

11. Page AJ, De Silva N, Hunt M, Quail MA, Parkhill J, Harris SR, et al. Robust high-throughput prokaryote de novo assembly and improvement pipeline for Illumina data. Microb Genom. 2016;2(8):e000083.

12. Holt KE, Parkhill J, Mazzoni CJ, Roumagnac P, Weill FX, Goodhead I, et al. High-throughput sequencing provides insights into genome variation and evolution in Salmonella Typhi. Nat Genet. 2008;40(8):987–93.

13. Ingle DJ, Nair S, Hartman H, Ashton PM, Dyson ZA, Day M, et al. Informal genomic surveillance of regional distribution of Salmonella Typhi genotypes and antimicrobial resistance via returning travellers. PLoS Negl Trop Dis. 2019;13(9):e0007620.

14. Croucher NJ, Page AJ, Connor TR, Delaney AJ, Keane JA, Bentley SD, et al. Rapid phylogenetic analysis of large samples of recombinant bacterial whole genome sequences using Gubbins. Nucleic Acids Res. 2015;43(3):e15.

15. Stamatakis A. RAxML version 8: a tool for phylogenetic analysis and post-analysis of large phylogenies. Bioinformatics. 2014;30(9):1312–3.

16. Page AJ, Cummins CA, Hunt M, Wong VK, Reuter S, Holden MT, et al. Roary: rapid large-scale prokaryote pan genome analysis. Bioinformatics. 2015;31(22):3691–3.

17. Page AJ, Taylor B, Delaney AJ, Soares J, Seemann T, Keane JA, et al. SNP-sites: rapid efficient extraction of SNPs from multi-FASTA alignments. Microb Genom. 2016;2(4):e000056.

18. Yoshida CE, Kruczkiewicz P, Laing CR, Lingohr EJ, Gannon VP, Nash JH, et al. The Salmonella In Silico Typing Resource (SISTR): An Open Web-Accessible Tool for Rapidly Typing and Subtyping Draft Salmonella Genome Assemblies. PLoS One. 2016;11(1):e0147101.

19. Zhou Z, Alikhan NF, Mohamed K, Fan Y, Agama Study G, Achtman M. The EnteroBase user’s guide, with case studies on Salmonella transmissions, Yersinia pestis phylogeny, and Escherichia core genomic diversity. Genome Res. 2020;30(1):138–52.

20. Argimon S, Yeats CA, Goater RJ, Abudahab K, Taylor B, Underwood A, et al. A global resource for genomic predictions of antimicrobial resistance and surveillance of Salmonella Typhi at pathogenwatch. Nat Commun. 2021;12(1):2879.

21. Wong VK, Baker S, Connor TR, Pickard D, Page AJ, Dave J, et al. An extended genotyping framework for Salmonella enterica serovar Typhi, the cause of human typhoid. Nat Commun. 2016;7:12827.

22. Hunt M, Mather AE, Sanchez-Buso L, Page AJ, Parkhill J, Keane JA, et al. ARIBA: rapid antimicrobial resistance genotyping directly from sequencing reads. Microb Genom. 2017;3(10):e000131.

23. Zankari E, Hasman H, Cosentino S, Vestergaard M, Rasmussen S, Lund O, et al. Identification of acquired antimicrobial resistance genes. J Antimicrob Chemoth. 2012;67(11):2640–4.

24. Zankari E, Allesoe R, Joensen KG, Cavaco LM, Lund O, Aarestrup FM. PointFinder: a novel web tool for WGS-based detection of antimicrobial resistance associated with chromosomal point mutations in bacterial pathogens. J Antimicrob Chemother. 2017;72(10):2764–8.

25. Jacoby GA. AmpC beta-lactamases. Clin Microbiol Rev. 2009;22(1): 161–82, Table of Contents.

26. Banerji S, Simon S, Tille A, Fruth A, Flieger A. Genome-based Salmonella serotyping as the new gold standard. Sci Rep. 2020;10(1):4333.

27. Santos PDM, Widmer KW, Rivera WL. PCR-based detection and serovar identification of Salmonella in retail meat collected from wet markets in Metro Manila, Philippines. PLoS One. 2020;15(9):e0239457.

28. Hopkins KL, Kirchner M, Guerra B, Granier SA, Lucarelli C, Porrero MC, et al. Multiresistant Salmonella enterica serovar 4,[5],12:i:-in Europe: a new pandemic strain? Euro Surveill. 2010;15(22):19580.

29. Mather AE, Phuong TLT, Gao Y, Clare S, Mukhopadhyay S, Goulding DA, et al. New Variant of Multidrug-Resistant Salmonella enterica Serovar Typhimurium Associated with Invasive Disease in Immunocompromised Patients in Vietnam. mBio. 2018;9(5).

30. Allard MW, Luo Y, Strain E, Pettengill J, Timme R, Wang C, et al. On the evolutionary history, population genetics and diversity among isolates of Salmonella Enteritidis PFGE pattern JEGX01.0004. PLoS One. 2013;8(1):e55254.

31. Chiou CS, Hong YP, Liao YS, Wang YW, Tu YH, Chen BH, et al. New Multidrug-Resistant Salmonella enterica Serovar Anatum Clone, Taiwan, 2015-2017. Emerg Infect Dis. 2019;25(1):144–7.

32. Hendriksen RS, Leekitcharoenphon P, Mikoleit M, Jensen JD, Kaas RS, Roer L, et al. Genomic dissection of travel-associated extended-spectrum-beta-lactamase-producing Salmonella enterica serovar typhi isolates originating from the Philippines: a one-off occurrence or a threat to effective treatment of typhoid fever? J Clin Microbiol. 2015;53(2):677–80.

33. Ashton PM, Nair S, Peters TM, Bale JA, Powell DG, Painset A, et al. Identification of Salmonella for public health surveillance using whole genome sequencing. PeerJ. 2016;4:e1752.

34. Deng X, den Bakker HC, Hendriksen RS. Genomic Epidemiology: Whole-Genome-Sequencing-Powered Surveillance and Outbreak Investigation of Foodborne Bacterial Pathogens. Annu Rev Food Sci Technol. 2016;7:353–74.

35. Franz E, Gras LM, Dallman T. Significance of whole genome sequencing for surveillance, source attribution and microbial risk assessment of foodborne pathogens. Current Opinion in Food Science. 2016;8:74–9.

36. Nadon C, Van Walle I, Gerner-Smidt P, Campos J, Chinen I, Concepcion-Acevedo J, et al. PulseNet International: Vision for the implementation of whole genome sequencing (WGS) for global food-borne disease surveillance. Euro Surveill. 2017;22(23).

